# Human electroencephalography recordings for 1,854 concepts presented in rapid serial visual presentation streams

**DOI:** 10.1101/2021.06.03.447008

**Authors:** Tijl Grootswagers, Ivy Zhou, Amanda K. Robinson, Martin N. Hebart, Thomas A. Carlson

## Abstract

The neural basis of object recognition and semantic knowledge has been extensively studied but the high dimensionality of object space makes it challenging to develop overarching theories on how the brain organises object knowledge. To help understand how the brain allows us to recognise, categorise, and represent objects and object categories, there is a growing interest in using large-scale image databases for neuroimaging experiments. In the current paper, we present THINGS-EEG, a dataset containing human electroencephalography responses from 50 subjects to 1,854 object concepts and 22,248 images in the THINGS stimulus set, a manually curated and high-quality image database that was specifically designed for studying human vision. The THINGS-EEG dataset provides neuroimaging recordings to a systematic collection of objects and concepts and can therefore support a wide array of research to understand visual object processing in the human brain.

## Background & Summary

Humans are able to visually recognise and meaningfully interact with a large number of different objects, despite drastic changes in retinal projection, lighting or viewing angle, and the objects being positioned in cluttered visual environments. Object recognition and semantic knowledge, our ability to make sense of the objects around us, have been the subject of a large amount of cognitive neuroscience research^1–3^. However, previous neuroimaging research in this field has often relied on a manual selection of a small set of images^1,4,5^. In contrast, recent developments in computer vision have produced very large image sets for training artificial intelligence, but the individual images in these sets are minimally curated and therefore make them often unsuitable for research in psychology and neuroscience. To overcome these issues, recent work has created large, curated image sets that are designed for studying the cognitive and neural basis of human vision^4,6^. One of these is THINGS^4^, which is an image set containing 1,854 object concepts representing a comprehensive set of nameable concepts used in the English language, accompanied by 26,107 associated manually-curated high-quality image exemplars and human behavioural annotations. This rich collection of stimuli and behavioural data has already been used to study the core representational dimensions underlying human similarity judgements^7^. The next phase is to collate corresponding neural responses to stimuli in THINGS. This would contribute to an emerging landscape of large datasets of neural responses to curated image sets that accelerate research in visual, computational, and cognitive neuroscience^8^.

Collecting neurophysiological data for the THINGS dataset, with over 26,000 images, is unachievable in a traditional neuroimaging experiment: Typically, classic object vision experiments present around one image per second ^e.g., 9–11^. Collecting one trial for each image in THINGS would take more than seven hours, which is infeasible to achieve with a single session and would require a complex design involving multiple scanning sessions. However, we have recently shown that it is possible to uncover detailed information about visual stimuli presented in rapid serial visual presentation (RSVP) streams using electroencephalography (EEG)^12–15^. In one of these studies^12^, participants viewed over 16,000 visual object presentations at 5 and 20 images per second, in a single 40-minute EEG session. Results from multivariate pattern classification and representational similarity analysis revealed detailed temporal dynamics of object processing that were similar to work that used slower presentation speeds (around one image per second). Therefore, fast presentation paradigms are highly suitable for collecting neural responses to the large number of visual object stimuli in the THINGS database.

Here, we present THINGS-EEG, a dataset of human (n=50) electro-encephalography responses to 22248 images from all 1,854 concepts in the THINGS object database. In the main session, each image was repeated once, and in the second part, 200 validation images were repeated 12 times each to be able to assess data quality and compare the data to future datasets acquired with other modalities. In total, 26,248 visual images were presented in a 1-hour EEG session. This was achieved using a rapid serial visual presentation paradigm. Technical validation results indicated the dataset contains detailed neural responses to images, which shows that the dataset is a high-quality resource for future investigations into the neurobiology of visual object recognition.

In this study, we presented over 25,000 trials in a 1-hour EEG experiment. While this is an exceptionally large number in a visual object perception study, the paradigm has several limitations. Firstly, by presenting the images in rapid succession at 10Hz, new information is being presented while previous trials are still being processed. While our previous work has shown that a great amount of detail about objects can be extracted from brain responses to RSVP streams^12–15^, the images are being forward and backward masked, and therefore the data does not capture the full brain response to each image^13^. For example, cognitive functions such as memories or emotions may not have enough time to be instantiated at such rapid presentation rates. Our design also involved one presentation per image, which makes image-specific analyses challenging, placing the focus of this work at the level of the 1,854 object concepts. The benefit of this is that the data has a built-in control for image-level confounds. For example, visual regularities that are specific to an image but vary across a concept will not generalise to the concept level. Visual statistics that reliably differ at the concept level of course may need to be accounted for, depending on the goals of the experimenter. For example, recent work has pointed out concept-level differences in mean luminance between images in the THINGS image set^16^, which can be controlled for in future analyses of the THINGS-EEG dataset. Another point to consider is that our recording setup did not include EOG or EMG channels, which means the dataset does not contain external recordings of eye or other muscle movements. These movement patterns are unlikely to contain informative stimulus-specific information, due to the fast presentation paradigm. However, they still cause noise artefacts in the EEG data. Future users could consider detecting and correcting for eye movements using the frontal EEG channels.

The THINGS-EEG dataset has strong potential for investigating the neurobiology of visual object recognition and semantic knowledge. The dataset presents a very large set of non-invasive neural responses to visual stimuli in human participants. We foresee many possible uses of this dataset. For example, the dataset could be used to test models of visual object representation, such as different semantic models, or deep neural networks. It could be used to test the generalisability of previous studies that were limited by small stimulus sets^5^. The rapid presentation paradigm allows to examine sequential effects in the data, such as how a specific object concept influences the encoding of the subsequent presentations. The consistent presentation frequency also lends itself to separate the data into oscillatory components. Instead of the RSA and classification analyses presented here, it is also possible to analyse the data in an encoding framework, for example by creating an encoding model from the semantic information in the THINGS-dataset. In sum, as THINGS is a high-quality stimulus set of record size, THINGS-EEG accompanies this resource with a comprehensive set of human neuroimaging recordings.

## Methods

50 individuals volunteered to take part in the experiment in return for course credit. Participants were recruited from the undergraduate student population at the University of Sydney. They were 36 females and 14 males, mean age 20.44 (sd 2.72), age range 17 – 30. Participants had different language profiles, with 26 native English speakers, 24 non-native speakers, 24 monolinguals, and 25 bilinguals. Handedness was not recorded. All participants reported normal or corrected-to-normal vision and reported no neurological or psychiatric disorders. There are 4 participants marked for potential exclusion due to notably poor signal quality or equipment failure (marked in the *participants*.*tsv* file). These participants are included in the release for completeness. The study was approved by the University of Sydney ethics committee. Informed consent was obtained from all participants at the start of the experiment.

Stimuli were obtained from the THINGS database^4^ (Figure 1A). For detailed information on the contents and organisation of this stimulus database, readers are referred to the accompanying publication^4^. THINGS contains 1,854 objects concepts, with 12 or more images per concept. The first 12 images for each concept were used for this experiment, resulting in 22,248 different visual images, which we divided into 72 sequences of 309 stimuli. Individual concepts were never repeated within one sequence. To be able to assess within- and between-subject variance on the same images, we presented another 200 validation images from the THINGS database 12 times to every subject after the main part of the experiment. For every subject we used the same 200 validation images (listed in the *test_images*.*csv* file). We repeated these images in 12 sequences of 200, where we presented them in random order. The experiment thus contained 84 sequences in total. The subjects were not explicitly made aware of the two different parts.

**Figure 1:**
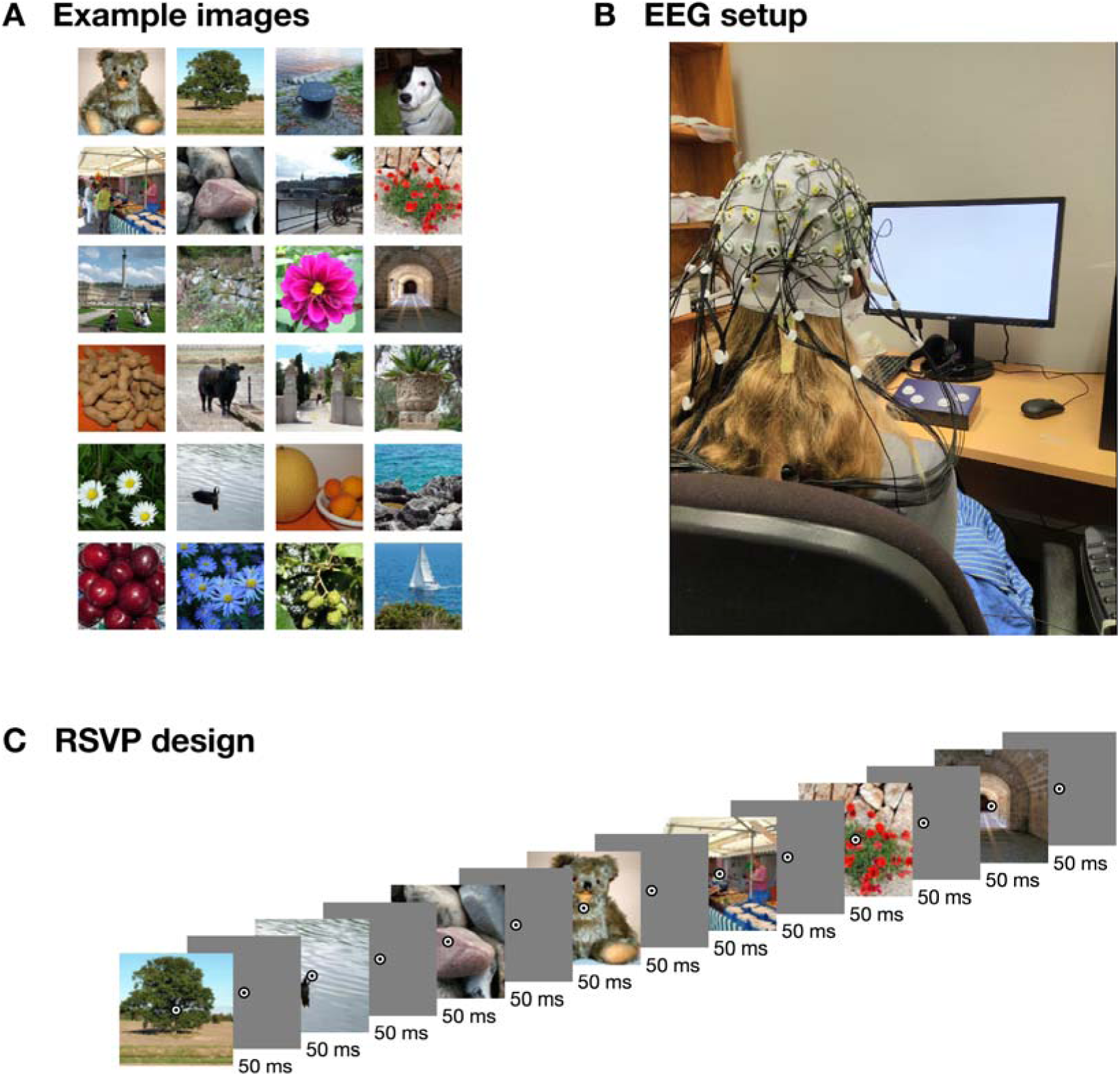
Example stimuli, design, and EEG setup image. A) Example images similar to the stimuli used in the experiment. B) EEG experimental setup (photo credit AKR). C) Rapid serial visual presentation design. For illustration purposes, only part of the sequence is shown. For this figure, all images were replaced by public domain images with similar appearance (obtained from PublicDomainPictures.net: Brunhilde Reinig).

The experiment was programmed in Python (v3.7), using the Psychopy^17^ library (version 3.0.5). The sequences were presented at 10Hz, with a 50% duty cycle (Figure 1C). That is, each image was presented for 50ms, followed by a 50ms blank screen. Participants were seated about 57cm from the screen, and the stimuli subtended approximately 10 degrees visual angle. Overlaid at the centre of each image was a bullseye (0.5 degrees visual angle) to help participants maintain fixation. To increase attention and engagement, each sequence contained 2 to 5 random target events, where the bullseye turned red for 100ms, and the participants were instructed to press a button on a button box using their right hand. These target events are marked in the dataset, as researchers may want to consider excluding these target events, depending on the aim of their analysis. At the end of each sequence, the display showed the progress through the experiment, and participants were able to start the next sequence with a button press. Participants were asked to sit still and minimise eye movements during the sequences and to use the time between sequences as breaks and relax, and start the next sequence using a button press when they were ready. The experiment lasted around one hour.

We used a BrainVision ActiChamp system to record continuous data while participants viewed the sequences (Figure 1B). Conductive gel was used to reduce impedance at each electrode site below 10 kOhm where possible. The median electrode impedance was under 18 kOhm in 40/50 participants and under 60 kOhm in all participants. We used 64 electrodes, arranged according to the international standard 10–10 system for electrode placement^18,19^. The signal was digitised at a 1000-Hz sample rate with a resolution of 0.0488281µV. Electrodes were referenced online to Cz. An event trigger was sent over the parallel port at the start of each sequence (trigger code E3), and at every stimulus onset event (trigger code E1) and stimulus offset event (trigger code E2).

To perform basic quality checks and technical validation, for each subject, we ran a standard decoding analysis. We decoded pairwise images for the 200 validation images, and we created the full time-varying 1,854×1,854 Representational Dissimilarity Matrix^20,21^ reflecting the pairwise decoding accuracies between all 1,854 object concepts. We used a minimal preprocessing pipeline derived from our previous RSVP-MVPA studies^12–15^. Using Matlab (R2020b) and the EEGlab (v14.0.0b) toolbox^22^, data were filtered using a Hamming windowed FIR filter with 0.1Hz highpass and 100Hz lowpass filters, re-referenced to the average reference, and downsampled to 250Hz. Epochs were created for each individual stimulus presentation ranging from [-100 to 1000ms] relative to stimulus onset. No further preprocessing steps were applied for the technical validation analysis presented here (as in our previous work using similar presentation paradigms^12–15^). Researchers may want to consider popular preprocessing steps such as baseline correction or eye movement correction. The channel voltages at each time point served as input to the decoding analysis.

Decoding analyses were performed in Matlab using the CoSMoMVPA toolbox^23^. We first decoded between the 200 validation images. For a given pairs of images, we used a leave-one-sequence out (total: 12 sequences) cross-validation procedure and trained a regularised (λ=0.01) linear discriminant classifier to distinguish between the images. The mean classification accuracy on the image pair in the left-out sequences were stored in a 200×200×275×50 (image×image×time point×subject) RDM, which is symmetrical across its first diagonal. A similar procedure was performed for the main experiment, using the 1,854 different image concepts. This resulted in an 1,854×1,854×275×50 (concept×concept×time point×subject) RDM. For the 200 validation images, we also computed noise ceilings by comparing between subject RDMs, as described in previous work^24^. The noise ceilings estimate the lower and upper bound of the highest achievable performance of a model that attempts to explain variance in the data.

### Data Records

All data and code are publicly available. The raw EEG recordings are hosted in BIDS^25,26^ format on OpenNeuro (https://doi.org/10.18112/openneuro.ds003825.v1.1.0)^27^. The preprocessed (Matlab/EEGLAB^22^ format) data, and group-average RDMs are included for convenience (in the data/derivatives directory). The RDMs for individual subjects are hosted in Matlab format in a separate repository on Figshare (https://doi.org/10.6084/m9.figshare.14721282)^28^. All custom code is available from the Open Science Framework (https://doi.org/10.17605/OSF.IO/HD6ZK)^29^, which also contains links to the above repositories.

### Technical Validation

We computed representational dissimilarity matrices for the 200 validation images, by calculating time-varying decoding accuracy between all pairs of images. Mean, subject-wise decoding accuracy (Figure 2A) showed an initial peak around 100ms, and a second, lower peak around 200ms after stimulus onset. The shape of the time-varying decoding was similar to previous object decoding studies^e.g., 9,10^, and was also similar to previous results on images presented in fast succession^12–14^, indicating data quality was similar to these studies. For most subjects, this shape was apparent from their individual data (Figure 2B). The noise ceiling (Figure 2C) indicates an average similarity (correlation) of up to 0.2 between the subject-specific dissimilarity matrices.

**Figure 2:**
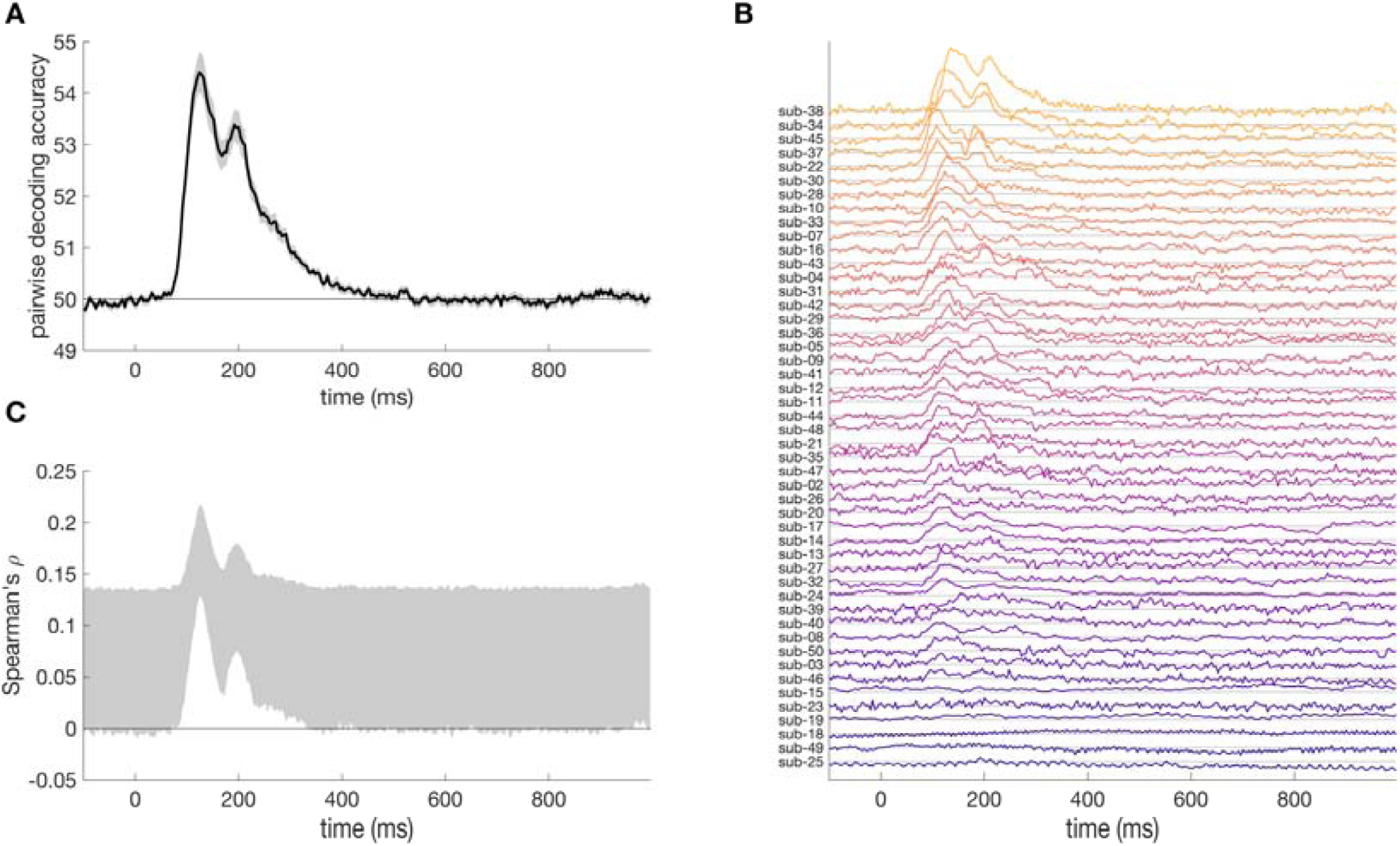
Results for the 200 validation images that were repeated 12 times at the end of the session. A) mean pairwise classification accuracy over time. B) Mean pairwise decoding over time, per subject, sorted by peak classification accuracy. Subjects 1 and 6 are not shown as they did not have data on the validation images. C) Noise ceiling over time shows the expected correlation of the ‘true’ model with the RDMs of the validation images and reflects the between-subject variance in the RDMs.

Next, we computed the full RDM for all 1,854 images concepts (1,717,731 pairs). The average accuracy within this RDM (Figure 3A) was lower than for the validation images, which is likely due to the fact that accuracy reflects generalised concept-similarity across images. Figure 3B shows the full RDM at one time point (200ms). To test if the values in the RDM contain meaningful information, we computed the correlation between the full RDM and four example categorical models (Figure 3C). The models coded for the presence of a certain category (e.g., animal). Figure 3C shows each model reaches an above-zero correlation, with the ‘natural’ model reaching the highest correlation, around 200ms.

**Figure 3:**
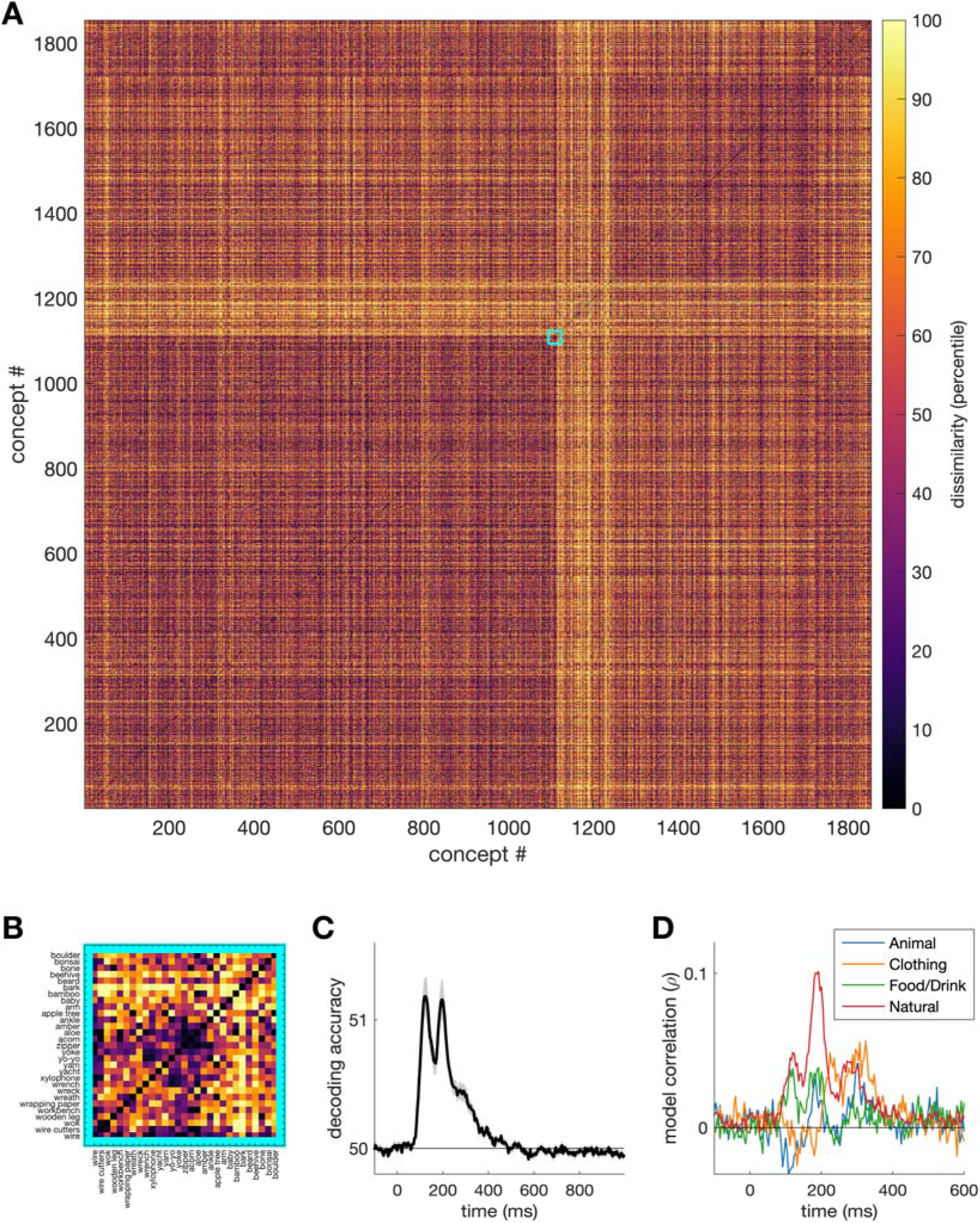
Results for the 1,854 image concepts that were repeated 12 times (using a different image each time). A) Full 1,854×1,854 RDM at 200ms, arranged by high-level category. B) Zoomed in section of the full RDM. C) Mean pairwise classification accuracy between concepts over time. D) Correlation over time between the neural RDM and four high-level categorical models.

## Code Availability

Code and detailed instructions to reproduce the technical validation analyses and figures presented in this manuscript are available from the Open Science Framework (https://doi.org/10.17605/OSF.IO/HD6ZK)^29^, which also contains links to the data repositories.

## Acknowledgements

This research was supported by ARC DP160101300 (TAC), ARC DP200101787 (TAC), and ARC DE200101159 (AKR). The authors acknowledge the University of Sydney HPC service for providing High Performance Computing resources.

## Author contributions

TG: Conceptualization, Methodology, Investigation, Formal analysis, Visualization, Data Curation, Writing – Original Draft, Writing – Review & Editing, Supervision, Project administration.

IZ: Conceptualization, Investigation, Data Curation, Writing – Review & Editing, Project administration.

AKR: Conceptualization, Methodology, Writing – Review & Editing.

MNH: Conceptualization, Methodology, Writing – Review & Editing.

TAC: Conceptualization, Methodology, Writing – Review & Editing, Supervision, Funding acquisition, Project administration.

## Competing interests

The authors declare no competing interests.

## Notes

https://osf.io/hd6zk/

### Competing Interest Statement

The authors have declared no competing interest.

